# Building a story: coherent narrative formation relies on functional connectivity in posterior cortex and frontoparietal networks

**DOI:** 10.1101/667618

**Authors:** Amir Assouline, Avi Mendelsohn

**Affiliations:** Sagol Department of Neurobiology, University of Haifa, Haifa 3498838, Israel; The Institute of Information Processing and Decision Making (IIPDM), University of Haifa, Haifa, Israel

## Abstract

Narratives are embedded in human experience, enabling the integration and communication of large quantities of accumulated information. Despite the ubiquity of this deeply rooted ability, the neural networks involved in narrative formation are yet unclear. Building on literary and philosophical definitions of narrative, we explored brain networks that differentially coactivated while individuals were presented with either coherent or incoherent narratives. Using movie scenes presented in a functional MRI environment, either in their correct or reversed order, we found that regions in the posterior cortex and frontoparietal networks were preferentially co-activated during coherent narrative formation. Moreover, whereas coactivation patterns of posterior cortex converged across conditions over time, the frontoparietal network remained constantly higher in the coherent narrative condition. We suggest that processing and integrating accumulating information is supported by functional coupling of posterior cortical networks, whereas the frontoparietal network serves to maintain the coherence and causal relations that underpin plot comprehension.

## Introduction

Narratives, or stories, are ancient concepts that emerged at the early days of human culture. From fables painted on cave-walls, through written stories and religious tells, a variety of genres for narrative communication have existed from ancient periods to common day culture. Some literary theorists and philosophers point to Aristotle’s ‘Poetic’ as the first document to delineate the basic definition of narrative in the west^1^. Narratives are thought to play an important role in the psychology of the self, as the ability to produce a narrative is pivotal to one’s sense of personal and cultural identity, and in the creation and construction of memories^2,3^. Certain psycho-pathologies such as schizophrenia are accompanied, among other things, by a disability to produce a coherent narrative of one’s current and past self or to compose a cohesive narrative from recalled events^4^. Moreover, narrative processing seems to be afflicted in post-traumatic stress disorder (PTSD)^5^, wherein coherent and organized accounts of the past trauma increase the likelihood of salutary gains^1,6,7^. Despite the essential role that narratives play in fundamental aspects of human experience, the neural mechanisms that support the formation of a succinct cohesive story, and particularly in binding related events that unfold over time, are not well characterized.

The concept of narrative, and particularly the associations that arise by this concept, are complex and therefore call for a clear definition. Typical narratives often portray a protagonist, who holds a certain goal (e.g. the purchase of a car) and whose progress towards that goal is impeded (e.g. the loss of a job) or facilitated ^1^. Aristotle, in his Poetics, emphasized the importance of coherence in narrative, which is the result of logically causal connections between occurrences ^8^. Here we rely on fundamental literary accounts of narrative, defined as a series of actions and events that unfold over time according to causal principles ^1^. To be considered a coherent narrative, events must occur in a logical order, such that the actions and episodes that lead to other events must take temporal precedence given the conflation of logical (if x then y), causal (because x then y), and temporal priority (first x then y) ^1,8,9^.

Successful construction of a coherent narrative from an episodic occurrence entails the interactive participation of several cognitive faculties, among them perceptual processing, working memory, social cognition and reasoning. Executive functions, and particularly working memory, are particularly imperative in the formation of coherent narratives by placing incoming occurrences against the backdrop of accumulated information pertinent to the story^10,11^. In narratives involving social encounters, intentions, and interactions among protagonists, social cognition processes such as theory of mind, empathy, and social inference of others are bound to take place during the process ^12–14^. Lastly, understanding the causal chain of events for forming a narrative necessitate^12^s high-order cognitive functions that enable the understanding of cause and effect.

It has been suggested that narrative formation provides the building blocks of episodic memory construction ^3^. Indeed, rather than scrambled flashes of past occurrences, episodic memory is organized as casually connected sequential sections that together compose a coherent story ^3,15,16^. Conversely, memories of traumatic events are characterized by incoherent contextually detached and temporally distorted representations ^5,7,16^. Recent findings have demonstrated that initial stages of consolidation begin immediately following the encoding of meaningful information that is suggested to bind together to form a compressed story ^12,17^. Thus, the time periods following such ‘event boundaries’ of information streams ^18,19^, characteristic of real-life occurrences or movie contents that propagate over time, are imperative in binding meaningful information into long-term representations. Real-life episodes, however, pose a challenge, as natural events typically unfold continuously, precluding the opportunity to investigate post-encoding processes that occur in conjunction with the flow of new incoming information. In the current study, we set out to explore both online and offline processing of continuous narrative formation using discrete film segments.

How does the brain form narratives? Previous studies have shown that the construction of short narratives from written sentences relies on regions in the medial prefrontal and parietal cortices ^19^. Transitions between information units in stories (i.e. narrative shifts) are reported to correlate with activations in the precuneus and posterior cingulate cortex^20^, as well as angular gyrus^21^, regions suggested to support event segmentation ^18^. In a series of studies using movies and stories, Hasson and colleagues found that large-scale high-order regions were activated both during viewing and recollecting narrative scenes ^22^, assigning a particular important role to the default mode network (DMN), whose activation similarity across participants corresponded to retrieval of similar narrative representations ^23^, and in which inter-subject correlations increased as the narrative unfolded. High order associative brain regions are particularly sensitive to narrative processing^24^. The superior temporal gyrus and inferior frontal gyrus respond similarly to narratives conveyed both by speech and visual input, showing indifference to input modality ^25^. A network of frontal, temporal and cingulate areas that support working-memory and theory-of-mind processes seem to be a recurring theme in narrative comprehension ^1^. Despite these and other advances in the field, the neural networks that support coherent narrative formation and particularly how they evolve during the course of narrative formulation are largely unknown. The temporal aspect of coherent narrative construction is of particular interest in this study, which aims at exploring whether and how the cohesiveness of narrative-related brain regions changes throughout the course of narrative building. The process of narrative formation entails the ongoing acquisition of new information and embedding it within previous relevant information units. Our working framework considers two main trajectories of narrative-related cognitive faculties that should play a part in narrative construction, enabling 1) the processing of time-dependent accumulation of information, and 2) the continuous representation of narrative-related information throughout. Conducting functional connectivity analysis enabled us to explore how networks of distributed brain regions co-activate during the unfolding of coherent vs. incoherent narratives.

We presented participants with scenes from the classic film ‘The Bicycle Thief’ (Directed by Vittorio de Sica, 1948), either in their correct order, conforming with the applied definition of narrative, or in an order that departs from the definition. In brief, the film follows the story of a man, who attains a bicycle as a condition to hold a new job. His bicycle is stolen at the outset, a dramatic event that sets the main protagonist on an excruciating path to retain them, upon which he himself ultimately deteriorates into attempting bicycle theft, only to fail and endure a humiliating experience witnessed by his young child. Members of the “incoherent narrative” group were presented with the same scenes, presented in reversed order, thus breaking temporal order and causal connection between events, impeding the formation of a coherent narrative according to the definition of narrative formation ^1,9^. By supplying the two groups with identical information items, yet manipulating essential components that influence coherent narrative formation, we explore neural correlates of coherent narrative formation. Focusing on functional connectivity measurements within and across groups, we dissociate brain networks that differentiate states of narrative formation from scene perception.

## Results

Participants were scanned in an fMRI environment while viewing movie scenes that were presented either in their chronological order (coherent group) or in reversed order (incoherent group), thus leaving in place or omitting a fundamental feature of narrative coherence, i.e. the causal connection between events (Fig. 1). The bulk of participants in the incoherent narrative group indeed noticed that the scenes were out of order, which coincided with their narrative understanding, as derived from their written description of the story, which tended to be unclear and ambiguous compared to the coherent narrative group. This was particularly evident by the participants’ responses to a question about the causal connection between two dramatically loaded subsequent scenes. This question was answered correctly by 78 percent of the coherent narrative group, compared to only 11 percent of the incoherent narrative group. In addition, when subjects were asked to describe the general plot in their own words, the members of the incoherent group were less successful in describing fundamental components of the plot, including the events’ order, the main protagonist’s goal and the plot’s development. This was despite the fact that the questionnaire was administered several minutes after the experiment, thus allowing retroactive narrative formation. Notably, neither the subjective affect during the movie (as rated in the post-movie questionnaire) nor the willingness to participate in future follow-up research differed between the groups.

**Figure 1.**
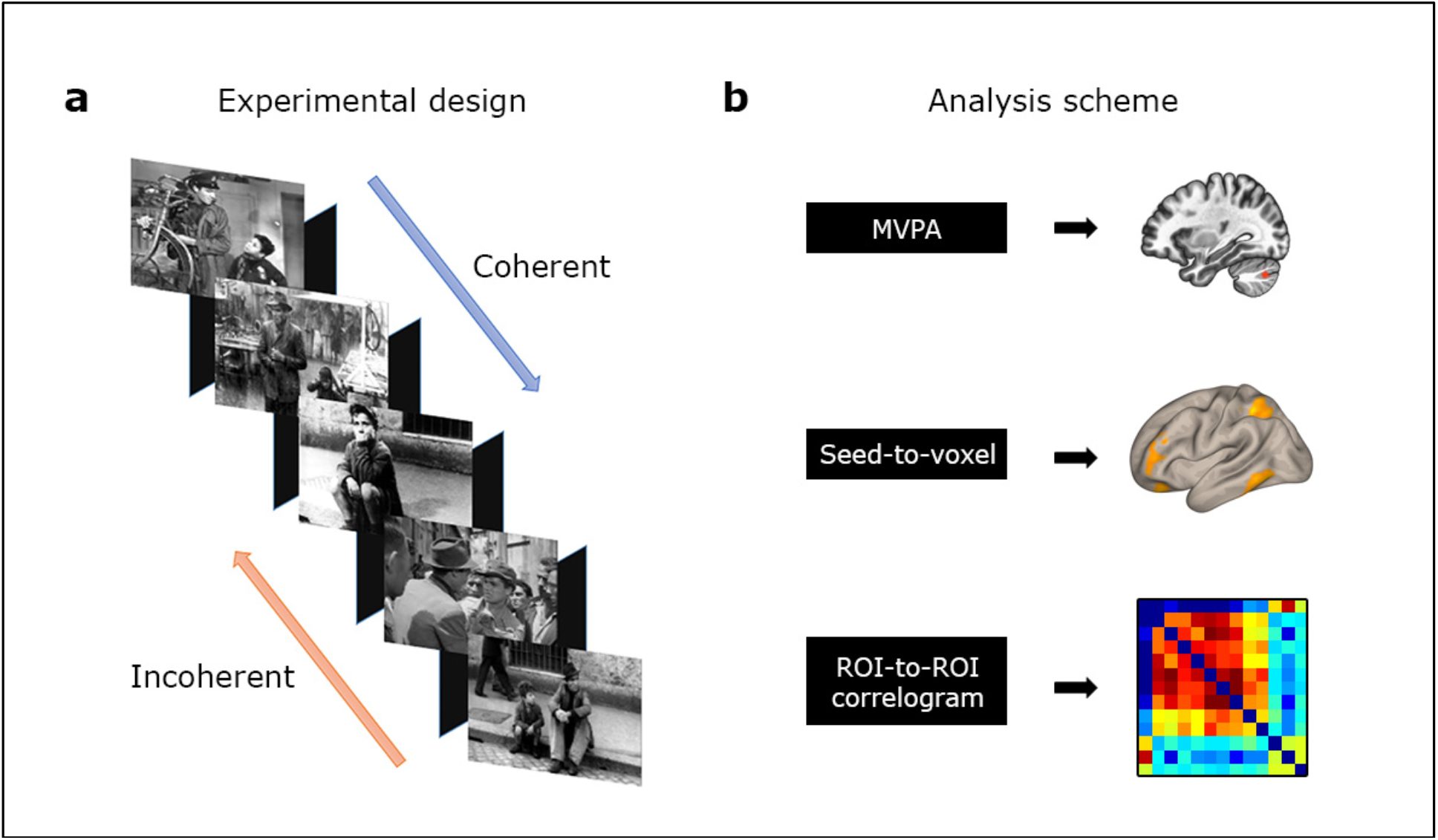
Experimental design and analysis outline. **(a)** Twenty-two scenes from the film ‘The Bicycle thieves’ were presented during the experiment (each lasting ~60sec), interleaved by blank screens (10 sec). The arrows indicate the direction of scene order as presented to the coherent (from beginning to end) and incoherent (from end to beginning) groups. **(b)** Multivariate pattern connectivity analysis (MVPA) analysis was performed to delineate regions that differed in functional connectivity between the two groups. This was followed by extracting regions of interest (ROIs) that served for seed-to-voxel analysis within and between groups, culminating in two distinct networks, which were subsequently depicted in correlograms and examined across time (see Methods and Result sections).

To explore networks that differed between the two groups, we performed a multivariate pattern analysis (MVPA) followed by seed-to-voxel analysis as described in the Methods section. After analyzing the clusters resulting from the MVPA map of the inter-scene ‘blank’ periods, we extracted two regions that significantly differed between the coherent vs. incoherent narrative groups (p<0.001, cluster level p≤0.05 false discovery rate (FDR) corrected). These regions consisted of the intra and supra calcarine (4,−88,12) and the cerebellum Crus II (−32,−74,−38). In the second stage of analysis, the two abovementioned ROIs served as origins of a seed-to-voxel analysis that yielded two distinct networks that were the result of a between-group analysis, referred to here as the calcarine-posterior cortex network (based on seed 4,−88,12) and the cerebellum-frontoparietal network (based on seed −32,−74, −38).

*The calcarine-posterior cortex network* was further divided into two distinct subnetworks, comprising regions that were positively or negatively correlated with the seed. The positive sub-network included the posterior precuneus and bilateral posterior hippocampus (*P*<0.001, cluster level threshold p<0.05, FDR corrected). The negative sub-network included clusters in the occipital pole, fusiform gyrus, lingual gyrus, bilateral superior parietal lobule, superior and middle frontal gyrus, (overlapping with Brodmann area 10) and the superior precuneus. Inspecting each group separately indicated that the negative sub-network (i.e., regions with lower functional connectivity in the coherent compared to incoherent group) stemmed from negative functional connectivity values with the seed region in the coherent group (Fig. 2a). The differences between the group were apparent both during blank and movie segments of the movie, yet were more pronounced in the blank periods (Fig. 2b).

**Figure 2.**
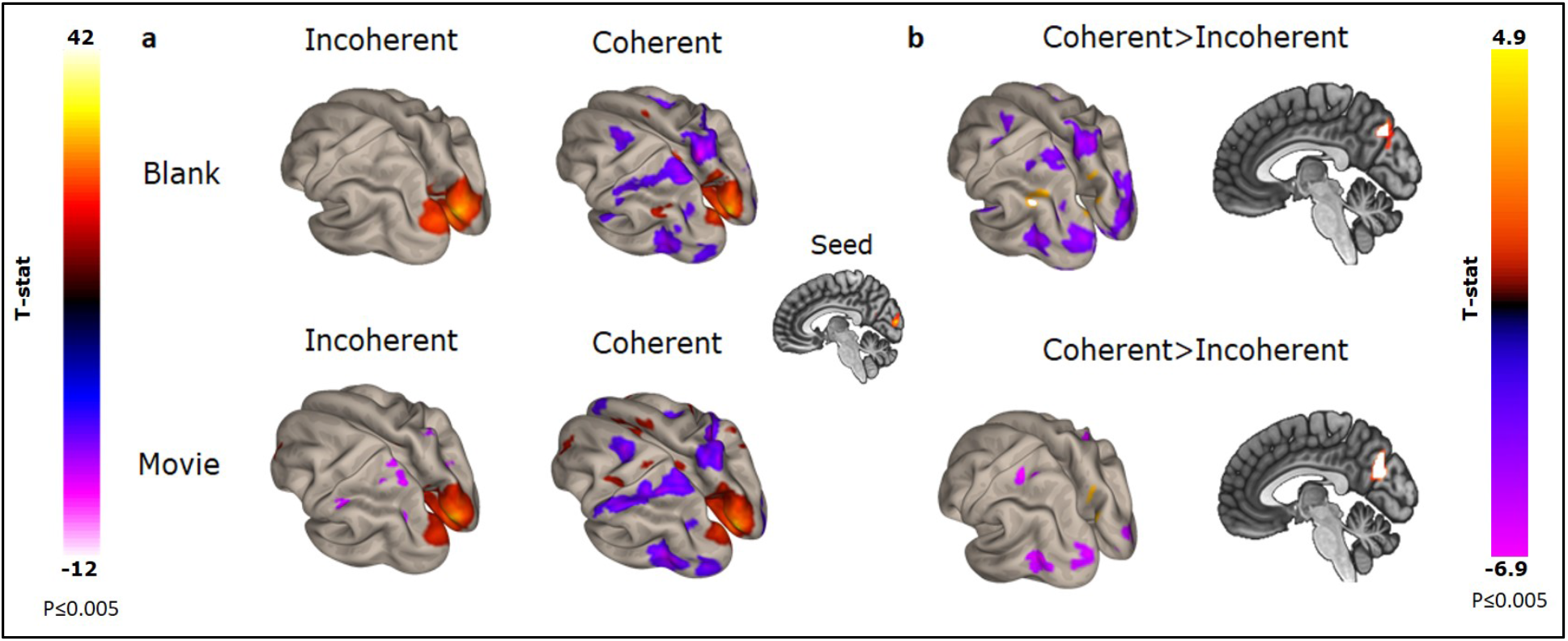
Seed-to-voxel analysis with calcarine as the seed. The seed region (MNI −4, −88, 12) is presented in the middle of the figure. **a.** Areas surpassing the applied statistical threshold of the seed-to-voxel whole-brain functional connectivity analysis is shown for each group separately and for each condition (Blank, top; Movie, bottom). Positive co-activations are shown in yellow-red hues, and negative in blue-purple. **b.** Direct between-group comparisons of functional connectivity with the calcarine as the seed. Enhanced functional connectivity for coherent > incoherent group is shown in red-yellow hues, whereas incoherent > coherent indices are colored with blue-purple. High-order sensory regions show higher functional connectivity with the seed region in the incoherent vs. coherent group (areas outlined in purple) both during blank and movie epochs. The precuneus and the angular (during the blank only) are correlated with the seed region to a higher degree in the coherent vs. incoherent group (top right panel).

The *cerebellum-frontoparietal network* exhibited positive correlations between the left cerebellum Crus II and extensive parts of the frontoparietal network in the coherent compared to the non-coherent narrative group. This network encompasses the inferior frontal gyrus, left orbital gyrus, bilateral supramarginal gyrus, middle frontal gyrus, inferior temporal gyrus, angular gyrus and superior parietal lobule (Fig. 3). These between-group differences in coactivation were more pronounced during the movie scenes than during blank epochs.

**Figure. 3.**
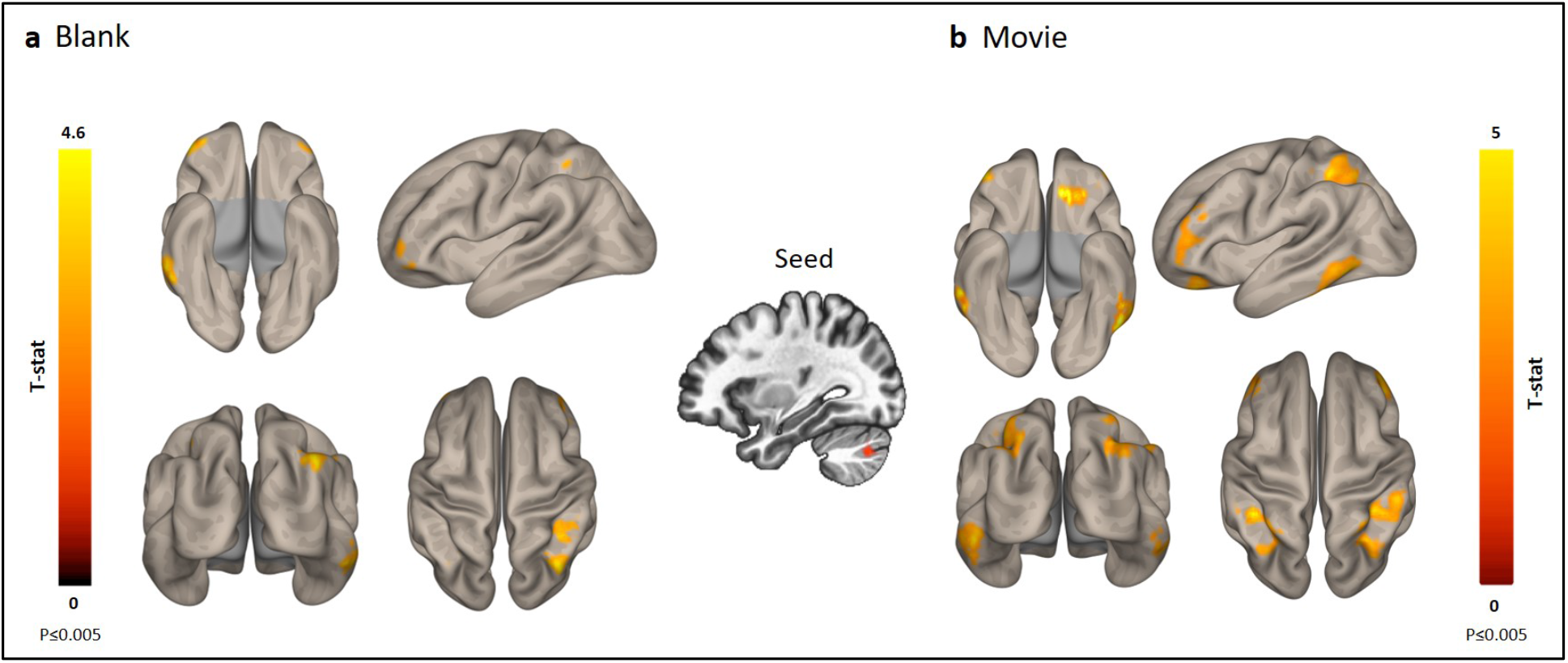
Seed-to-voxel analysis with the cerebellum Crus II as a seed. The seed ROI (MNI −32 −74 −38) is presented in the middle of the figure. Areas surpassing the applied statistical threshold of the seed-to-voxel whole-brain functional connectivity analysis are shown for Blank epochs (**a**) and Movie sections (**b**). Differences in functional connectivity with the seed region between the groups are observed in parietal, frontal and temporal lobes, largely overlapping the frontoparietal network. The differences are more pronounced during the movie (**b**) than during blank segments (**a**).

### Temporal dynamics of coherent narrative formation

To explore the intrinsic functional connectivity within the networks reported above along the experimental time-line, we computed correlations between the ROIs of each network for the beginning, middle and end of the experiment, plotted ROI correlograms for each group (see Supplementary Material) and depicted the mean differences between the groups in matrices (Fig. 4a & 5a). In the *calcarine-posterior cortex network*, the differences in ROI-to-ROI correlations between the coherent and incoherent group are particularly pronounced in the first segment of the experiment (top left matrix plot), followed by decreases in the differences with time. Although the differences between the two groups in the seed-to-ROI correlations are maintained throughout the experiment (as seen in the first row and column of the matrices), the group-differences within and between the networks weaken with time (Fig. 4a).

**Figure 4.**
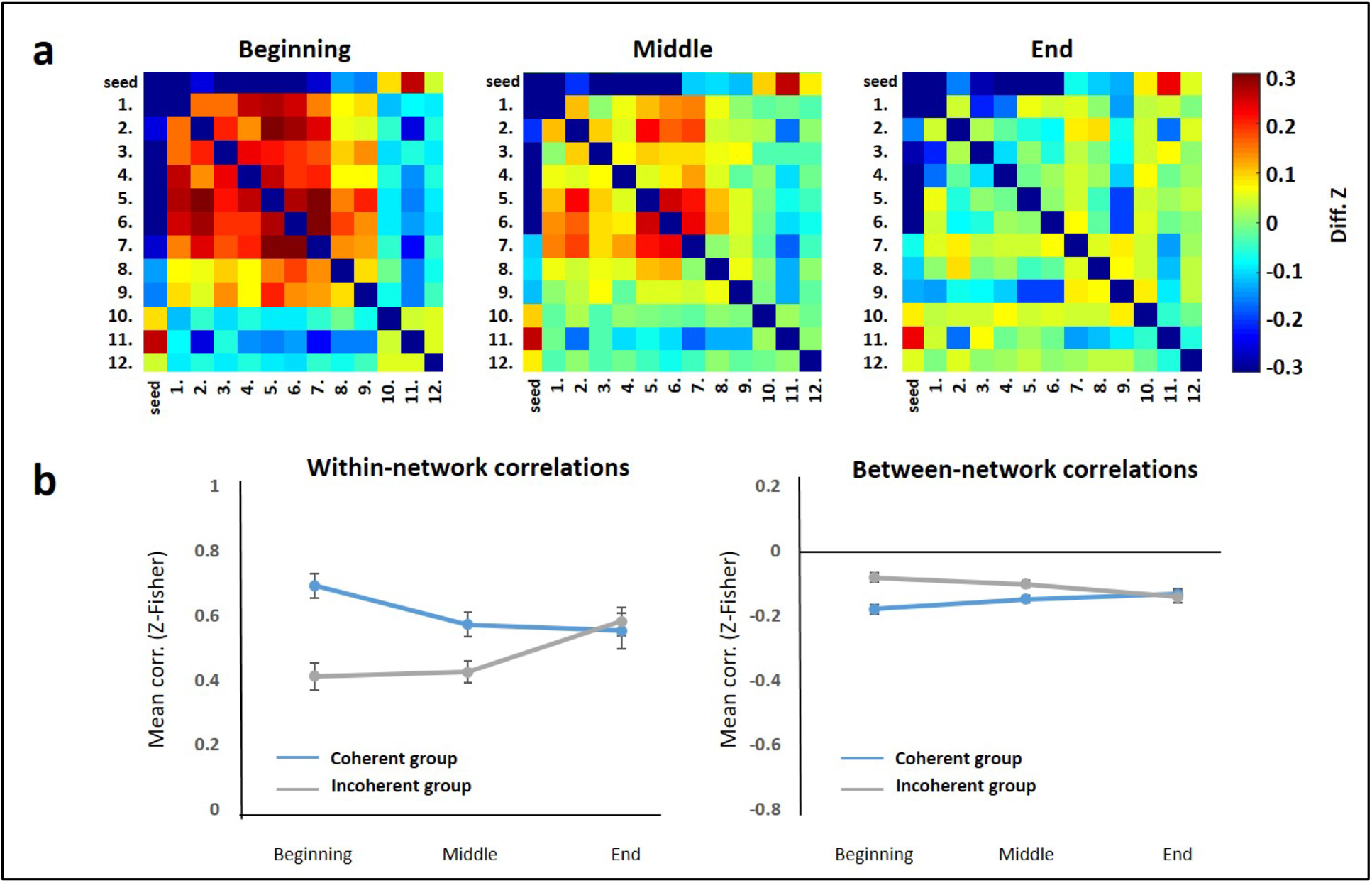
Between-group differences in ROI-to-ROI correlations across time. **a.** Matrices depicting differences in ROI-ROI correlations of the calcarine-precuneus network between the coherent and incoherent groups during the beginning, middle, and end of the experimental timeline (see Table 1 for ROI details). **b.** Mean correlations within the seed-to-ROI networks (regions 1-9 and 10-12 among themselves, left panel), and between the two networks (correlations between regions 1-9 and 10-12, right panel) for beginning, middle, and end of the experiment. Within-network correlations differed between groups in the beginning (*P*<0.0001) and middle sections of the movie (*P*<0.0001), but not in the final third (*P*=0.59), yielding a main effect for group (F_(1,36)_=6.248, *P*<0.05) and a group by time interaction effect (F_(2,72)_=21.92, *P*<0.0001). Similarly, between network correlations significantly differed between the groups in the beginning (*P*<0.0001) and middle sections (*P*<0.05), but not in the final third (*P*=0.71), yielding a main effect for group (F_(2,36)_=13.03, *P*<0.05) and an interaction effect (F_(2,72)_=16.25, *P*<0.0001).

**Table 1.**
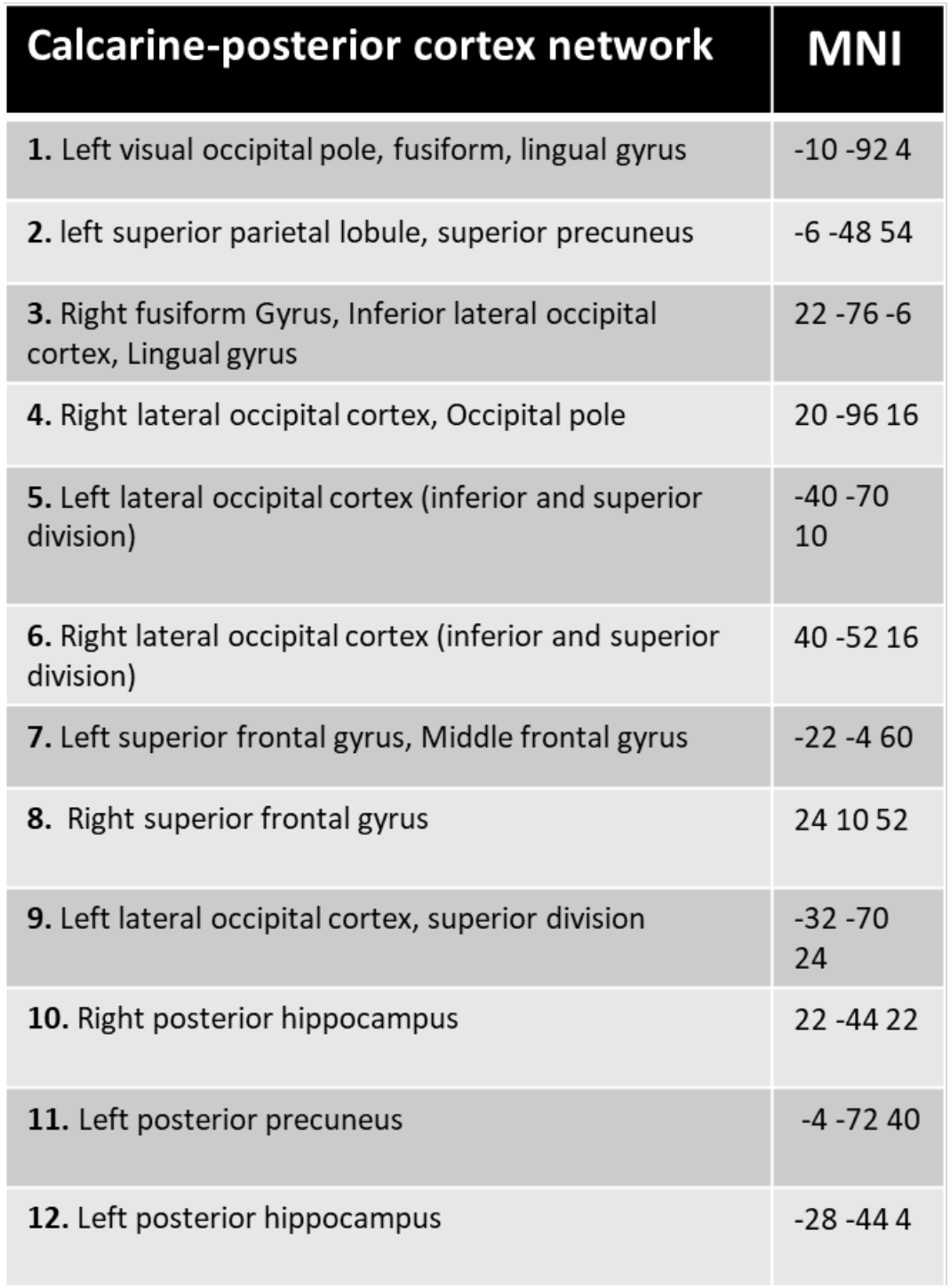
Regions of the calcarine-posterior cortex network. Listed regions correspond to the list of numbered regions depicted in the correlograms of Fig. 4a. The MNI coordinates indicate voxels of peak significance.

| Calcarine-posterior cortex network | MNI |
| --- | --- |
| 1. Left visual occipital pole, fusiform, lingual gyrus | -10 -92 4 |
| 2. left superior parietal lobule, superior precuneus | -6 -48 54 |
| 3. Right fusiform Gyrus, Inferior lateral occipital cortex, Lingual gyrus | 22 -76 -6 |
| 4. Right lateral occipital cortex, Occipital pole | 20 -96 16 |
| 5. Left lateral occipital cortex (inferior and superior division) | -40 -70 10 |
| 6. Right lateral occipital cortex (inferior and superior division) | 40 -52 16 |
| 7. Left superior frontal gyrus, Middle frontal gyrus | -22 -4 60 |
| 8. Right superior frontal gyrus | 24 10 52 |
| 9. Left lateral occipital cortex, superior division | -32 -70 24 |
| 10. Right posterior hippocampus | 22 -44 22 |
| 11. Left posterior precuneus | -4 -72 40 |
| 12. Left posterior hippocampus | -28 -44 4 |

This notion is corroborated by analysis of within- and between-network correlations across time, revealing that within-network correlations were more positive in the coherent compared to the incoherent group at the beginning of the experiment, and converge as time unfolded (main effect of group: F_(1,36)_=6.248, *P*<0.05; interaction of group by time: F_(2,72)_=21.92, *P*<0.0001; Fig. 4a left panel). Similarly, between-network correlations (Fig. 4b, right panel) were more negative in the coherent narrative group compared to the incoherent group at the outset, and again converged over time (main effect of group F_(1,36)_=13.033, *P*<0.05; interaction of group by time: F_(2,72)_=16.25, *P*<0.0001). These correlation patterns may indicate that the initial story segments are the ones most critical to the construction of a coherent narrative.

For the regions comprising the cerebellum-frontoparietal network, we performed the same analysis as for the calcarine network, i.e., subtracted the correlation matrix of incoherent narrative group from the coherent narrative group and repeated the process across the time span (Fig. 5a). In contrast to the calcarine-posterior cortex network, the group differences in correlations within the network remained constant across the experiment sections, as apparent both from the matrices and the line graphs (Fig. 5). Thus, throughout the three segmented sections of the experimental timeline, the mean ROI-to-ROI correlations were higher in the coherent vs. incoherent narrative group, leading to a main effect for group across the experiment sections (F_(1,36)_=7.039, *P*<0.05), and no main effect of time or group by time interaction (Fig. 5b).

**Figure 5.**
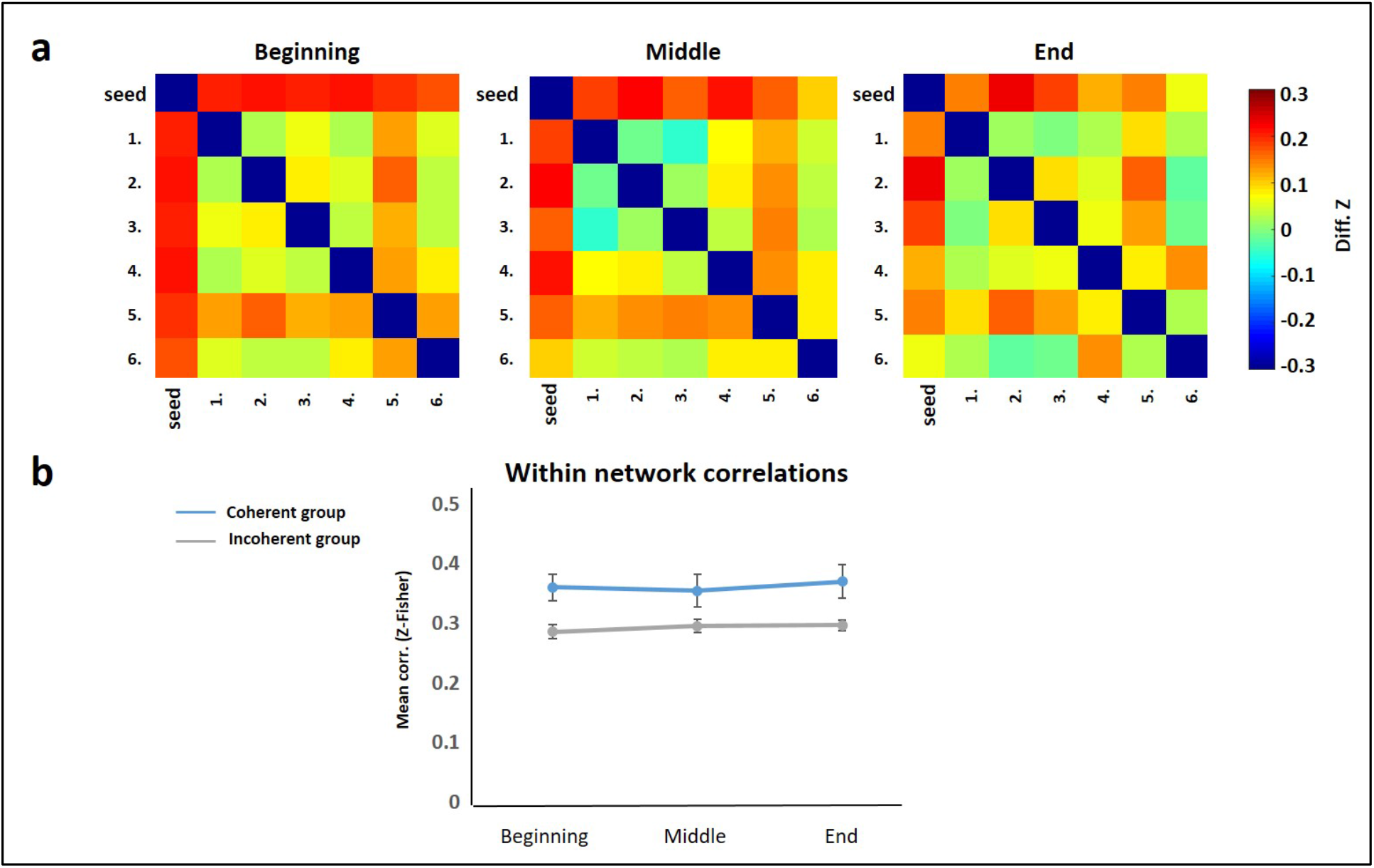
ROI-to-ROI co-activation analysis across the experiment. **a.** Matrices depicting ROI-ROI correlations within the cerebellum-frontoparietal network, which differed between the coherent and incoherent groups during the beginning, middle, and end of the experimental timeline (see Table 2 for ROI details). **b.** Mean correlation values (normalized to Z-Fisher scores) across all ROI pairs along the experimental timeline demonstrate higher ROI-to-ROI correlations in the coherent group for beginning (*P* < 0.05), middle (*P* = 0.068) and final third (*P* < 0.005), yielding a main effect for group (F_(1,36)_=7.039, *P* < 0.05) and no interaction (F_(2,72)_=0.373, *P*=0.69). Diff.: difference, corr.: correlation.

**Table 2.** Regions of the cerebellum-frontoparietal. Listed regions correspond to the list of numbered regions depicted in the correlograms of Fig. 5a. The MNI coordinates indicate voxels of peak significance.

| Cerebellum-frontoparietal network | MNI |
| --- | --- |
| 1. Right superior parietal lobule | 42 -46 40 |
| 2. Middle frontal gyrus | -4 48 46 |
| 3. Right inferior and middle temporal gyrus | 60 -48 -20 |
| 4. Left superior parietal lobule | -36 -32 56 |
| 5. Left cerebellum Crus II | -36 -78 -46 |
| 6. Left frontal pole | -36 60 12 |

Due to its pivotal role in episodic memory, and particularly in post-encoding ^17,28^ and event segmentation^19,26,27^, we investigated hippocampal functional connectivity patterns with the rest of the brain across the experimental timeline^28^. The results of this analysis were that the hippocampus exhibited functional connectivity primarily with components of the two networks identified above, and to a higher extent in the coherent compared to the incoherent group. In relation to time-dependent changes, initial segments of the film were characterized by stronger co-activations of hippocampus and precuneus in the coherent group, whereas in final segments of the movie (the third third), differential co-activations were detected between hippocampus and portions of the frontoparietal network (*P*≤0.001, cluster level *P*≤0.05 with FDR correction) (Fig. 6). Notably, the exhibited coherent > incoherent group hippocampus-precuneus co-activation during the first movie part derived from negative functional connectivity values in the incoherent group. The higher co-activations of hippocampus and frontoparietal regions in the final movie part is a consequence of steadily increasing functional connectivity of the hippocampus and the frontoparietal network in the coherent narrative group.

**Figure. 6.**
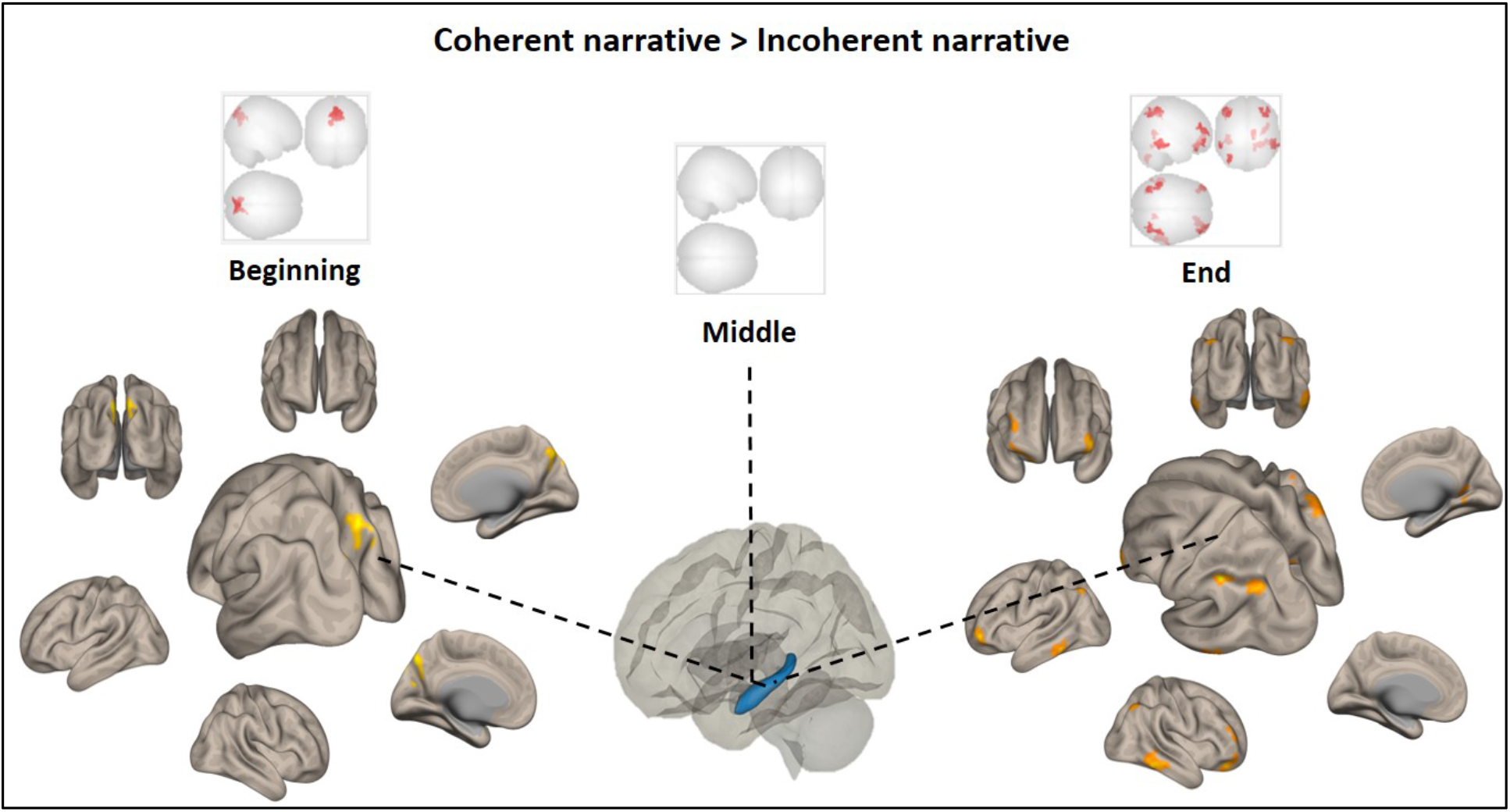
Whole-brain seed-to-voxel functional connectivity with hippocampus as seed. The left hippocampus (in blue at the center) served as a seed for a whole-brain seed-to-voxel analysis separately for beginning, middle and end of the experimental timeline. Statistical maps on inflated brain images indicate areas that were significantly different between the two groups (coherent > incoherent). In the first third (left), the hippocampus showed higher correlations with the precuneus in the coherent compared to the incoherent group (*P*<0.001). In the middle section of the movie, no significant differences were found between the groups, and in the final third, correlations between the hippocampus and parts of the frontoparietal network and inferior temporal gyrus were detected (*P*<0.001).

**Table 3.**
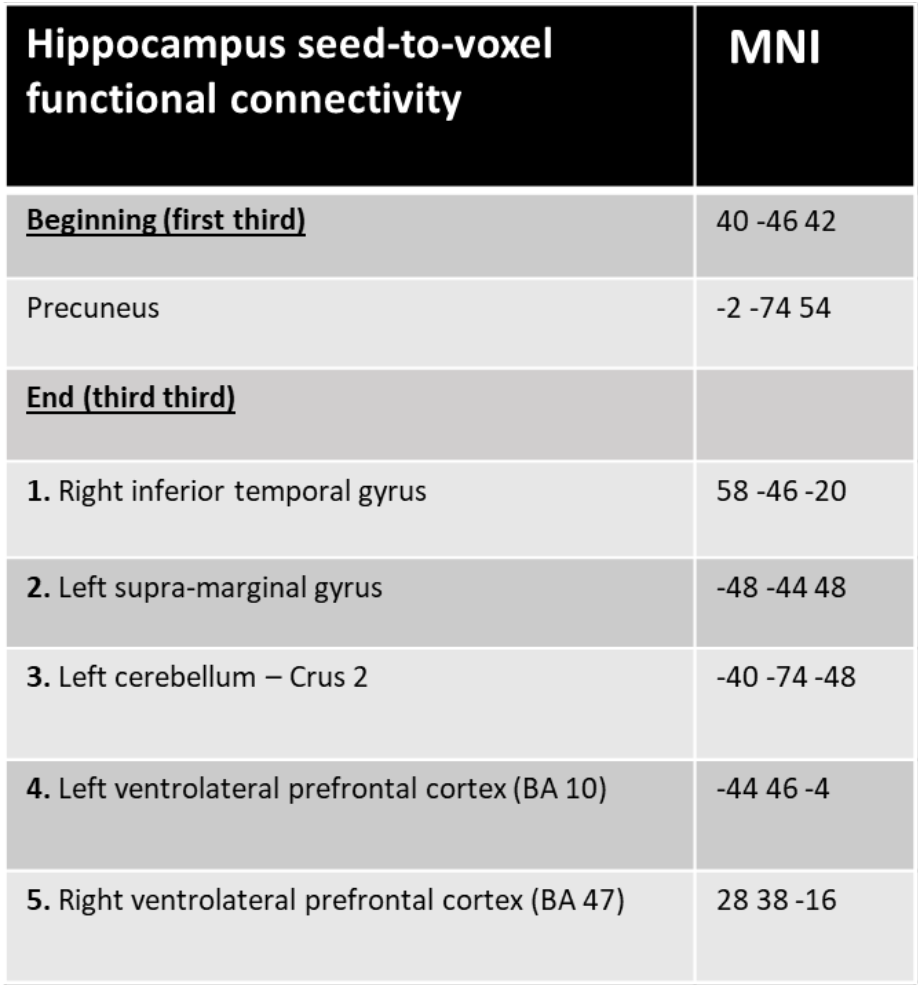
Regions resulting from seed-to-voxel analysis with hippocampus as seed. Listed regions correspond to the regions depicted in Fig. 6. MNI coordinates indicate voxels of peak significance.

| Hippocampus seed-to-voxel functional connectivity | MNI |
| --- | --- |
| <u>Beginning (first third)</u> | 40 -46 42 |
| Precuneus | -2 -74 54 |
| <u>End (third third)</u> |  |
| 1. Right inferior temporal gyrus | 58 -46 -20 |
| 2. Left supra-marginal gyrus | -48 -44 48 |
| 3. Left cerebellum – Crus 2 | -40 -74 -48 |
| 4. Left ventrolateral prefrontal cortex (BA 10) | -44 46 -4 |
| 5. Right ventrolateral prefrontal cortex (BA 47) | 28 38 -16 |

## Discussion

Narratives are of essential importance to human experience, playing a fundamental role in cognition and culture. Nevertheless, the neural mechanisms that underlie the formation and comprehension of narratives are still largely unknown, particularly as they pertain to binding accumulating information across time to form a coherent story. In the current study, we designed a novel experimental paradigm using an edited movie of the Italian Neorealism genre, which encapsulates core features of narrative characteristics in a cinematic medium ^1,29^. We provided at the outset the central criteria that are imperative to the construction of a coherent narrative, as well as exclusion criteria, i.e., what a narrative is not (a controversial issue in the literature field ^30^). This was based on both historical and literary accounts, as well as clinical considerations of healthy and pathological self-narrative ^31,32^, under the assumption that such definitions are deeply rooted in the basic psychological and cultural aspects of human nature^1^. As described above, we compared behavioral and neural responses to the same stimuli (cinematic scenes), which differed only by the presence or absence of the three most fundamental aspects of narrative: causality, directionality, and narrative coherence (as pointed by Aristotle, see Introduction). By eliminating the natural trajectory of content accumulation by reversing scene order, we sought to present the same occurrences, yet omit the core defining features that enable coherent narrative formation.

The behavioral findings indicate that although participants in both groups made an effort to provide an account of a perceived storyline, the incoherent narrative group failed to form a simple causal connection between two successive scenes that were a central theme of the narrative, even when they asserted a retroactive comprehension of the general story line. The fact that awareness of the reversed scene order was not enough to produce narrative coherence emphasizes the pivotal role that the original time axis has in causality inference ^33^, and hence in coherent narrative formation, substantiating the fundamental organizing principle of episodic memory, as introduced by Tulving^15^. The usage of naturalistic stimuli, namely continuous and meaningful audio-visual information in the form of films, has been gaining popularity in recent neuroimaging studies ^34^, particularly those exploring episodic memory^35,36^ and social cognition^12^. Expanding the ecological validity of cognitive tasks by shifting from reductionist approaches of lab-based tasks to naturalistic stimuli has contributed to the illumination of neural networks that underlie real-life cognition ^37,38^. Going beyond the abovementioned benefits of ecological validity, here we manipulated the core features of a coherent narrative, while maintaining in both cases the exact same stimuli. Moreover, the length of the movie was important in order to examine the temporal dynamics of network co-activation during the course of the movie’s unfolding. The usage of a prolonged set of scenes taken from a single movie offered the opportunity to investigate the temporal dynamics of coherent narrative formation over the course of information accumulation, and not only of the narrative as a whole.

Our analysis focused on delineating functional connectivity profiles devoid of prior assumptions regarding brain regions that may be differentially recruited during film observation in the coherent and incoherent narrative groups. This approach yielded two major networks characterized by discrete within- and between-network correlations across time, deemed to represent different functional aspects of coherent narrative representation. Previous studies on functional connectivity during narrative comprehension of auditory stories have detected coactivation of language-related regions with sensorimotor and dorsolateral prefrontal areas^39^ as well as with superior and middle temporal gyri ^40,41^. Notably, whereas these studies used auditory narration of stories, ours conveyed audio-visual information in the form of a foreign-language movie. Therefore, our results illuminate narrative processing networks beyond those that are dedicated to language comprehension and forming meaningful sentences, attributed to the essence of narrative formation - the binding of events that form a coherent and meaningful story representation.

The first network we detected consists of regions that were negatively or positively correlated with the calcarine seed and to a higher extent in the coherent compared to the incoherent narrative group, forming in effect two sub-networks. One of the regions found in this network was the precuneus, showing higher positive functional connectivity with the seed region in its posterior part, and negative correlations in its superior part in the coherent compared to the incoherent narrative group. The precuneus plays an important role in high cognitive functions^42^ and is involved in retrieval of past representations and in simulating future scenarios ^43^ as well as in meta-cognition^44^, all of which are critically required for coherent narrative construction. Its activation during the processing of a continuous story has previously been interpreted as being involved in the holistic representation of meaningful pieces of information ^45,46^ and their segmentation ^18,27,46^.

The angular gyrus (which showed increased correlations with the calcarine sulcus in the coherent narrative group) has a significant role in episodic memory^47^ and particularly in enabling the subjective experience of remembering ^48^. In addition, the precuneus and the angular gyrus are both part of the default mode network (DMN), the precuneus as a central hub^49^ and the angular as a key parietal node ^50,51^. The connection between episodic memory formation and narrative construction is a natural one, as can be inferred by the definition of narrative (see Introduction), which entails the binding of events in time and space^52^. A set of regions that highly co-activated in the coherent vs. incoherent group comprised of areas of the ventral and dorsal visual processing pathways, which integrate visual information for identification and action planning, respectively^53^. Specifically, the detected areas here include extrastriate occipital pole, fusiform gyrus, lingual gyrus, superior parietal lobule and the superior precuneus, collectively involved in high-order sensory functions, required for (visually portrayed) narrative formation. It is possible that the anti-correlation between these regions and the primary visual cortex, and only in the coherent narrative group, signifies a detachment of these high-order functions from the original sensory input, integrating incoming information as part of a contextual episode.

Both sub-networks (positive and negative differential correlations with the calcarine) demonstrate a change in co-activation pattern across time, such that the differences in the networks’ co-activation between the groups steadily declines as the movie unfolds, as within-network correlations converge in the final movie segments. This effect stems from a decline in within-network functional connectivity in the coherent narrative group and a parallel increase in the incoherent group (see Fig. 4b). These time-dependent changes in co-activation within the network of regions that play a functional role in high-order sensory processing, may indicate a pivotal involvement of these regions in narrative weaving, and specifically in forming a narrative time-line by embedding accumulative information into temporal context. This notion is consistent with the finding of relatively low network co-activation in the incoherent group at early stages of the movie, when scenes were less likely to be perceived as part of an integrated whole, resonating with previous work on information accumulation in high order cortical regions^54^. As information accumulated with scene progression, parallel to the networks’ gradual increase in cohesiveness, a broader contextual story line was established, enabling the construction of a narrative, be it accurate or not.

The second network found to differ between the coherent and non-coherent groups included regions that were co-activated with the Crus II area of the cerebellum, consisting largely of the frontoparietal network^55,56^. As in our study, the frontoparietal network has previously been shown to co-activate with the cerebellum’s Crus II ^56^, which has been implicated as part of a supramodal zone of the cerebellum ^57^. Serving as a key control system of the human brain, the frontoparietal network is considered to support information integration in various cognitive contexts ^58^, and was shown to play a role in context-dependent narrative comprehension ^59^. Resonating with the current finding, the frontoparietal was found to correlate with chronological vs. inconsistent information in a narrative comprehension task ^60^. The network’s correlation was construed as meeting working memory demands for integrating incoming and previous information. In the current study, the differences in this network’s functional connectivity between the groups remained stable during the unfolding of the movie. In addition, this network was more prominent during movie compared to blank segments, implying its involvement in online processing of incoming information. These two said features are contrasted with the characteristics of the calcarine-precuneus within-network correlations, which were more prominent during blank periods and converged over time between the groups. Taken together, we suggest that in the context of narrative construction, whereas the frontoparietal network seems to engage in online representation of previous and incoming information, the posterior-cortex network may be primarily involved in cumulative integration of information.

Adding to decades of research on its crucial role in declarative memory, recent studies have offered insights regarding the role that the hippocampus and surrounding cortices play in time perception and integration of accumulating information over time. Thus, hippocampus activation was shown to be involved in time perception^61 62^, in post-encoding of naturalistic film segments^17,26^, and event segmentation^19,26,27^. In addition, evidence from human and animal studies applying various physiological and behavioral approaches suggest that the hippocampus is essential to memory of the order of sequential events. Damage to the hippocampus impairs memory of the temporal order of events, and the hippocampus is activated in humans during encoding and recall of a sequence of events^61^. Here we report that the hippocampus exhibits differential group-related functional connectivity specifically with components of the two networks identified in our analysis, i.e. the posterior cortex and frontoparietal networks. Furthermore, as to time-dependent co-activation changes, differential hippocampus-precuneus functional connections were found in the beginning of the film presentation, whereas in final segments of the movie, differential hippocampal-frontoparietal co-activations were detected. Specifically, the hippocampal-precuneus effect here stemmed from negative correlations between these regions in the incoherent group. Negative functional connectivity between brain networks can delineate systems involved in competing processes^55^. For instance, negative correlations between the hippocampus and the dorsal attention network^62^ have been ascribed to their divergent functions in attending to internal mental states versus external stimuli, respectively^55^. Similarly, the negative correlation between the hippocampus and the precuneus may signify competing processes of these two regions in the context of the current task. It is therefore suggested here that the negative correlations in activation of hippocampus and precuneus may signify competing functions of processing and binding sequences of incoming stimuli on the one hand (mediated by hippocampus), and integrating accumulating information on the other (mediated by the precuneus)^42,49,61^. As such, when processing an ongoing narrative presented in its natural form, there is less of a mismatch between the present event and the progressing storyline^63,64^.

The enhancement in hippocampal-frontoparietal functional connectivity in the coherent-narrative group is in line with the frontoparietal network’s role in integrating working memory with attentional resource allocation^65^. It is also consistent with the role that the frontoparietal network, particularly in prefrontal and parietal cortex, play in temporally extended integration of information, by maintaining internal representations over sustained intervals^66^. These functions are pertinent to the process of plot formation, which demands the integration of incoming and previously accumulated information^61^. Our findings suggest that co-activation of the hippocampus and areas of the frontoparietal network tighten during the final stages of film presentation, suggesting that demands for information accumulation and integration, which are bound to increase as the narrative unfolds, is dependent, at least to some extent, on hippocampus-frontoparietal co-activation.

A unique aspect of the current work lies in the cross-discipline approach applied by assuming literary and philosophical definitions and ideas to the design and investigation of their neural bearings. Such an integrative approach, combined with an ecological experimental design, may lead to a deeper understanding of how the brain achieves the goal of information compression into relatively short stories that can be later retrieved and conveyed to others. Narratives are of great importance not only to normal human functionating but also to mental health, as evident by the observation that narrative formation is commonly impaired in various mental illnesses, particularly PTSD and Schizophrenia. A better understanding of the neural mechanisms that support narrative formation holds the potential for detection and interventions for those suffering from serious mental illnesses. In the field of episodic memory research, the current study provides new insights into the processing of information that unfold over relatively long durations of time (several minutes), unearthing wide-range cerebral co-activity that subserves narrative construction and plot comprehension.

## Methods

### Subjects

Thirty-eight healthy individuals (mean age 27 ± 3.7 y) participated in the study. Two separate groups of 19 subjects were assigned to the Coherent-Narrative or the Incoherent-Narrative groups (8 and 9 females respectively in the coherent and incoherent narrative groups). All participants had normal visual acuity (without glasses or corrective lenses, enabling eye tracking). None of the subjects were familiar with the movie presented during the scan. Experimental procedures were approved by a Helsinki Committee, as required by Israeli law. All subjects provided written informed consent prior to the experiment.

### Stimuli and Experimental Design

In the MRI scanner, participants watched either the coherent or incoherent narrative (experimental conditions), which consisted of 22 scenes (mean scene length: 55.15±23 sec) edited from the movie “The Bicycle Thief” (Italy, 1948), interleaved by 10 sec blank screens. The whole experiment lasted 25 minutes. The scenes were identical across groups and differed only in their order of presentation, such that in the coherent-narrative group, scenes were presented in the correct order-from first to last, and in the incoherent narrative scenes were presented in reverse order, from last scene to first. The movie was edited using “Movie-Maker” software (Microsoft ©), and were presented using Presentation Version 20.1 software (www.neurobs.com).

### Behavioral Assessment

Immediately following scanning, comprehension of the story was assessed using a questionnaire targeting narrative comprehension and personal ratings regarding the movie. Participants were asked, among other questions, whether the scenes were presented in their correct order, and if not, at which stage they had noticed that. They were also asked about the casual connection between two succeeding scenes in the middle of the movie in order to detect their understanding of the event’s causation, which is a central component in the definition of coherent narrative.

### MRI Data Acquisition and Preprocessing

Whole-brain imaging was performed in a 3T Siemens Magnetom MRI system (Siemens Medical Systems, Erlangen, Germany) using a 16-channel head coil. Blood-oxygenated-level-dependent (BOLD)-sensitive T2*-weighted functional images were acquired using a single shot gradient-echo EPI pulse sequence (TR = 2000 ms, TE = 30 ms, flip angle = 82°, 64 axial slices, 2mm^3^, FoV = 192 × 192 mm, interleaved slice ordering) and corrected online for head motion. Participants completed the task in two fMRI runs, each lasting ~12.5 minutes. The first two volumes were discarded to allow for equilibration effects. Visual stimuli were presented on a screen behind the scanner using Presentation software (www.neurobs.com), and were viewed through a mirror attached to the head coil. Following functional imaging, a high-resolution T1 scan was acquired for anatomic normalization.

Imaging data were slice-time corrected and realigned using the SPM12 package (Wellcome Institute of Cognitive Neurology, London). Functional volumes were co-registered, normalized to the MNI (Montreal Neurological Institute) template brain, and smoothed with an 8 mm^3^ isotropic Gaussian kernel. We assessed task-related functional connectivity using the CONN toolbox (http://www.nitrc.org/projects/conn) in MATLAB. The implemented CompCor routine was carried out for each participant, aimed at identifying principal components associated with white matter (WM) and cerebrospinal fluid (CSF), which were segmented^67^. These components, as well as realignment correction information were entered as covariates in the first-level model. Since CompCor accounts for the effects of subject movement, the global BOLD signal was not entered into the regression model.

### Multivariate pattern analysis

In order to detect brain networks that differed in functional connectivity patterns between the two groups, we performed a multivariate pattern analysis (MVPA). A key benefit of MVPA is the ability to differentiate between cognitive states based on activation patterns across voxels rather than in specific voxels, hence avoiding the information waste that derives from omitting sub-threshold voxels when those voxels can still contain meaningful information as part of a wider pattern ^34^. Another benefit of MVPA is its ability to characterize differences in functional connectivity patterns without presumptions concerning a-priori seed regions. We used the MVPA function embedded in the CONN toolbox, which includes reducing data dimensionality and using Eigenvectors to compute correlations between each voxel and the rest of the brain, followed by F-tests to trace voxels that demonstrate significantly different functional connectivity patterns between the groups. The significantly different voxels were then projected onto statistical maps that represent the differences in the functional activity between the coherent and incoherent narrative groups. We performed the MVPA analysis separately for the Blank and Movie segments. From the resulting MVPA maps, we extracted data from functional clusters in which the statistical significance of the Coherent > Incoherent group comparison was at a height threshold level of p<=0.001 and cluster threshold of p<0.05, FDR corrected. For visual depiction, we used a threshold of 0.005 (See figure legends).

Since MVPA is an omnibus test that informs on differences between groups but not on the source of differences^68^ (Abdi & Williams, 2010), further post-hoc exploration was carried out. Specifically, from the MVPA results, we delineated two seed regions-of-interest (ROIs) and used them to explore networks that differed between the two groups in their co-activation patterns. The seeds were situated in the calcarine sulcus and area Crus II of the cerebellum, from which first-level analysis was conducted, whereby correlation maps of each of the selected seeds with the rest of the brain were computed based on Fisher-transformed bivariate correlation coefficients between the seeds’ BOLD time-series (averaged across all voxels within each seed) and all other voxels. Following this, a second-level was carried out, where whole-brain between-group t-tests were carried out on the resulting Fisher’s z-transformed correlations. From the between-groups analysis we produced two separate statistical maps (one from each seed) that represented the difference between the groups and which served for analysis of the temporal dynamics (see below).

In order to assess the effects of time on the abovementioned networks, we divided each subject’s dataset into three segments of equal length, corresponding to beginning, middle and end of the experimental time-line. We then performed an ROI-to-ROI correlation analysis between the regions comprising each of the networks, aiming to characterize potential changes in co-activation within each of the two networks during the course of the experiment. For each time segment, we then calculated the average correlation score (after Z-Fisher transformation) for each pair of regions in each network, averaged the correlations across each group and produced a matrix graph that depicts the group differences. To assess the networks’ coherence across the three time points in each group, we further averaged the Z-Fisher correlation values for each subject across all pairs of regions within each network, and averaged this index across subjects of each group. For the calcarine-posterior cortex network, we also performed a similar analysis on the correlations between ROI pairs that showed opposing functional connectivity patterns with the seed region in the seed-to-voxel analysis (i.e., with the calcarine seed). This analysis was performed in order to assess the changes in integrity between the two subnetworks found in this analysis across time. We hypothesized that since the accumulation of information should serve to increase narrative coherence in both groups, the differences in network coherence initially seen between the two groups (as gauged by ROI-to-ROI correlations) would decline with time. Additionally, due to its significant role in temporal integration of episodic events, we conducted a seed to voxel analysis for each time-segment, using an atlas-defined anatomical delineation of the hippocampus as a seed.

